# A Gated Graph Transformer for Protein Complex Structure Quality Assessment and its Performance in CASP15

**DOI:** 10.1101/2022.05.19.492741

**Authors:** Xiao Chen, Alex Morehead, Jian Liu, Jianlin Cheng

## Abstract

**Motivation:** Proteins interact to form complexes to carry out essential biological functions. Computational methods such as AlphaFold-multimer have been developed to predict the quaternary structures of protein complexes. An important yet largely unsolved challenge in protein complex structure prediction is to accurately estimate the quality of predicted protein complex structures without any knowledge of the corresponding native structures. Such estimations can then be used to select high-quality predicted complex structures to facilitate biomedical research such as protein function analysis and drug discovery.

**Results:** In this work, we introduce a new gated neighborhood-modulating graph transformer to predict the quality of 3D protein complex structures. It incorporates node and edge gates within a graph transformer framework to control information flow during graph message passing. We trained, evaluated and tested the method (called DProQA) on newly-curated protein complex datasets before the 15th Critical Assessment of Techniques for Protein Structure Prediction (CASP15) and then blindly tested it in the 2022 CASP15 experiment. The method was ranked 3rd among the single-model quality assessment methods in CASP15 in terms of the ranking loss of TM-score on 36 complex targets. The rigorous internal and external experiments demonstrate that DProQA is effective in ranking protein complex structures.

**Availability:** The source code, data, and pre-trained models are available at https://github.com/jianlin-cheng/DProQA

**Supplementary information:** Supplementary data are available at *Bioinformatics* online.

## 1 Introduction

Proteins perform a broad range of biological functions. Protein-protein interactions (PPI) play a key role in many biological processes. Understanding the mechanisms and functions of PPIs may benefit many areas such as drug discovery (Scott *et al*. (2016); Athanasios *et al*. (2017); Macalino *et al*. (2018) and protein design Kortemme and Baker (2004); Baker (2006); Lippow and Tidor (2007)). Typically, high-resolution 3D structures of protein complexes can be determined using experimental solutions (e.g., X-ray crystallography and cryo-electron microscopy). However, due to the high costs associated with them, these methods cannot meet all the increasing demand of protein complex structures in modern biological research and technology development. In the context of this practical challenge, computational methods for protein complex structure prediction have recently been receiving an increasing amount of attention.

Recently, AlphaFold2-Multimer (Evans *et al*. (2021)), an end-to-end system for protein complex structure prediction system, improves the accuracy of predicting multimer (quaternary) structures considerably. However, compared to AlphaFold2’s outstanding performance for monomer (tertiary) structure prediction (Jumper *et al*. (2021)), the accuracy level for protein quaternary structure prediction still has much room for progress. One specific problem of predicting quaternary structures is the estimation of model accuracy (EMA) (also called quality assessment(QA), which plays a significant role in ranking and selecting predicted quaternary structural models of good quality (Kinch *et al*. (2021)).

However, unlike the tertiary structure quality assessment with many machine learning, particularly deep learning methods developed for evaluating the quality of tertiary structural models over many years, there are few deep learning methods for evaluating the quality of quaternary structural models. Existing EMA methods have not leveraged cutting-edge attention-based transformer-like deep learning architectures (Vaswani *et al*. (2017)) to enhance the quaternary structure quality assessment. The structural models in most existing datasets for training quaternary structure quality assessment methods (Liu *et al*. (2008); Lensink and Wodak (2014); Kotthoff *et al*. (2021)) were generated by traditional protein docking methods (Tovchigrechko and Vakser (2006); Pierce *et al*. (2011)), whose quality is much lower than the structural models predicted by the state-of-the-art protein complex structure predictors such as AlphaFold-multimer (Jumper *et al*. (2021); Bryant *et al*. (2022)). Consequently, EMA methods trained using these datasets may not work well on the structural models generated by the latest, more accurate protein complex structure predictors.

In general, EMA methods for predicted structures can be divided into two categories: multi-model methods and single-model methods. Multi-model methods take a pool of protein structural models as input and may use a comparison between the models to evaluate their quality, such as in the procedure performed by Pcons (Lundström *et al*. (2001)), ModFOLDClust (McGuffin (2007)), and DeepRank2 (Chen *et al*. (2021)). In contrast, single-model methods give a certain quality score for each protein structural model without considering other models’ information, such as in the procedure performed by ProQ2 (Uziela and Wallner (2016)), ProQ3 (Uziela *et al*. (2016)), DISTEMA (Chen and Cheng (2022)), and GNN_DOVE (Wang *et al*. (2021)). In this work, we develop a single-model Deep Protein Quality Assessment method (DProQA) for predicting the quality of protein complex structural models. In particular, DProQA introduces a Gated Graph Transformer, a novel graph neural network (GNN) that learns to modulate its input information to better guide its structure quality predictions. DProQA takes a single protein complex structure as input to predict its quality via a single forward pass. Moreover, it uses a multi-task learning strategy to predict the real-valued quality score of a structural model as well as classify it into multiple quality categories.

## 2 Related Work

### Protein tertiary and quaternary structure prediction

Predicting protein structures has been an essential problem for the last several decades. Recently, the problem of protein tertiary structure prediction has largely been solved by deep learning methods (e.g., Jumper *et al*. (2021)). Furthermore, new deep learning methods (e.g., Evans *et al*. (2021); Bryant *et al*. (2022); Guo *et al*. (2022)) have begun making advancements in protein complex (quaternary) structure prediction.

### Protein representation learning

Protein structures can be represented in various ways. Previously, proteins have been represented as tableau data in the form of hand-crafted features (Chen *et al*. (2020)). Along this line, many works (Wu *et al*. (2021); Chen and Cheng (2022)) have represented proteins using pairwise information embeddings such as residue-residue distance maps and contact maps. Recently, describing proteins as graphs has become a popular means of representing proteins, as such representations can learn and leverage proteins’ geometric information more naturally. For example, EnQA (Chen *et al*. (2023)) used 3D-equivariant graph representations to estimate the per-residue quality of protein structures. GVP (Jing *et al*. (2020)) uses directed Euclidean vectors to represent the positions of atoms of proteins for protein design and quality assessment tasks.

### Machine learning for protein structure quality assessment

Over the past few decades, various EMA methods for ranking and scoring protein complex structural models have been developed (Gray *et al*. (2003); Huang and Zou (2008); Vreven *et al*. (2011); Basu and Wallner (2016b); Geng *et al*. (2020)). Among these scoring methods, machine learning-based EMA methods have shown better performance than physics-based (Dominguez *et al*. (2003); Moal *et al*. (2013) and statistics-based methods Zhou and Zhou (2002); Pons *et al*. (2011)).

Recent machine learning methods have utilized various techniques and features to approach the task. For example, ProQDock (Basu and Wallner (2016b)) and iScore (Geng *et al*. (2020)) used protein structural features as the input for a support vector machine to predict model quality. EGCN (Cao and Shen (2020)) assembled graph pairs to represent protein complex structures and then employed a graph convolutional network (GCN) to learn graph structural information. DOVE (Wang *et al*. (2020)) used a 3D convolutional neural network (CNN) to extract features from protein-protein interfaces to predict model quality. In a similar spirit, GNN_DOVE (Wang *et al*. (2021)), PPDocking (Han *et al*. (2021)), and DeepRank-GNN (Réau *et al*. (2023)) trained Graph Attention Networks (Veličković *et al*. (2017)) to evaluate protein complex decoys. Moreover, PAUL (Eismann *et al*. (2021)) used a rotation-equivariant neural network to identify accurate models of protein complexes.

### Deep transformers

Increasingly more works have applied transformer-like architectures or multi-head attention (MHA) mechanisms to achieve state-of-the-art results in different domains. For example, the Swin-Transformer achieved state-of-the-art performance in various computer vision tasks (Hu *et al*. (2019); Liu *et al*. (2021, 2022)). Likewise, the MSA Transformer (Rao *et al*. (2021)) used tied row and column MHA to extract features from multiple sequence alignments of proteins. Moreover, DeepInteract (Morehead *et al*. (2021b)) introduced the Geometric Transformer to model protein chains as graphs for protein interface contact prediction.

### Contributions

Our work builds upon prior works by making the following contributions:

1. We provide the *first* example of applying transformer representation learning to the task of protein complex structure quality assessment, by introducing the new gated graph transformer architecture to iteratively update node and edge representations using the adaptive feature modulation.
2. The DProQA method was trained using the newly-developed protein complex datasets in which all structural decoys were generated using AlphaFold2 (Jumper *et al*. (2021)) and AlphaFold-Multimer (Evans *et al*. (2021)).
3. Using the newly-developed Docking Benchmark 5.5-AF2 (DMB55-AF2), we demonstrate the state-of-the-art performance of DProQA in comparison with the existing methods such as ZRANK2 (Pierce and Weng (2008)), GOAP (Zhou and Skolnick (2011)) and GNN_DOVE (Wang *et al*. (2021)).
4. DProQA was blindly tested in the 15th community-wide Critical Assessment of protein Structure Prediction (CASP15) in 2022, where it ranked 3rd among all single-model EMA methods in terms of ranking loss of structural models’ TM-score.

## 3 Methods and Materials

Illustrated from left to right in Figure 1, DProQA first receives a 3D protein complex structure as input and represents it as a K-NN graph. Notably, all chains in the complex are represented within the same graph, where pairs of atoms from the same chain are distinguished using a binary edge feature (i.e., in the same chain or not). Therefore, it can uniformly deal with a complex consisting of any number of chains. Moreover, it only requires a single protein complex structure as input without using any extra information such as multiple sequence alignments (MSAs) and residue-residue co-evolutionary features extracted from MSAs. Its output includes a real-valued quality score of the structure as well as a quality class it is assigned to.

**Fig. 1.**
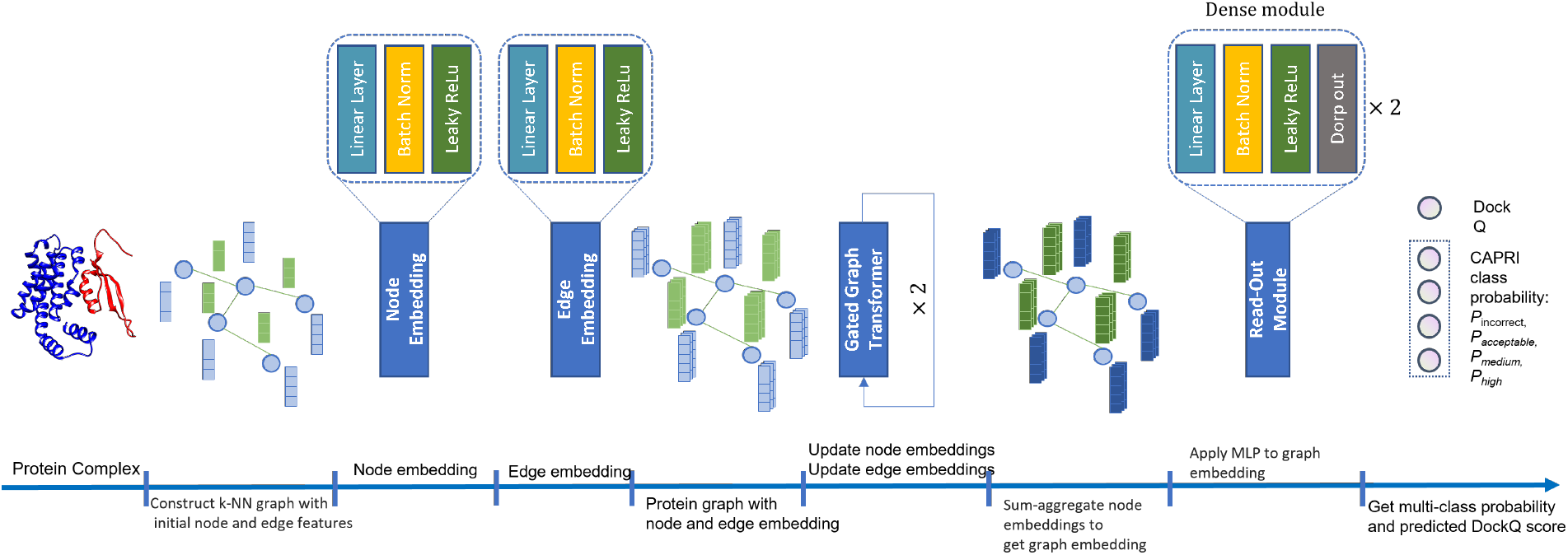
An overview of the DProQA pipeline for quality assessment of protein complexes. The input is a protein complex structure. The output includes the predicted quality score of the input structure (e.g., DockQ score) and the probability of the quality class (i.e., incorrect topology, acceptable quality, medium quality, and high quality) which the structure is classified into.

### 3.1 K-NN graph representation of protein complex structure

#### K-NN graph representation

K-nearest neighbors (K-NN) has been used in many previous studies for protein structure analysis and molecular representation learning (Ingraham *et al*. (2019)). Moreover, using other graph construction techniques (e.g. distance-based definitions of edges) may introduce inherent graph-structural biases (Jing *et al*. (2020)) within a network’s learning process. Subsequently, in this work, DproQA represents each input protein complex structure as a spatial k-nearest neighbors (k-NN) graph 𝒢 = (𝒱, *ε*), where the protein’s C*α* atoms serve as 𝒱 (i.e., the nodes of 𝒢). After constructing 𝒢 by connecting each node to its 10 closest neighbors in ℝ^3^, we denote its initial node features as **h** and its initial edge features as **e**.

#### Node and edge featurization

Table 1 describes DproQA’s node and edge features. Each node has 35 features and each edge has 6 features. For each graph 𝒢, the shape of node features is *N ×* 35, and the shape of the edge features is *E ×* 6, where *N* is the number of nodes and *E* is the number of edges.

**Table 1.**
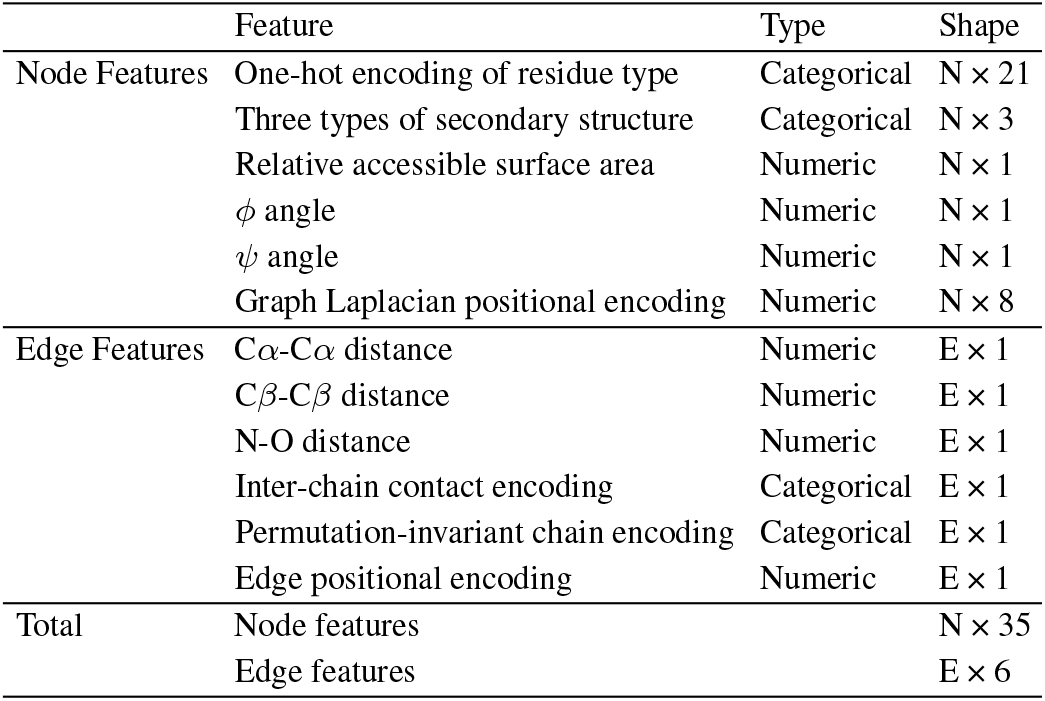
Summary of DProQA’s node and edge features. Here, N and E denote the number of nodes and edges in a protein graph, respectively.

The node features for a node include the one-hot encoding of 20 residue types as well as the 3-type secondary structures, relative solvent accessible surface area, and two torsion angles (*ϕ* and Ψ) computed by BioPython 1.79 (Cock *et al*. (2009)). The *ϕ* and Ψ angle values are normalized by the min-max normalization to scale their value range from [−180, 180] to [0, 1]. The graph Laplacian positional encoding (Dwivedi and Bresson (2020)) is added to each node.

The edge features for an edge include the distances between C*α* atoms, between C*β* atoms, and between backbone nitrogen and oxygen atoms of two residues. A binary feature indicating if two residues is in contact (i.e., if their C*α*-C*α* distance is less than 8Å) is added for each edge. To encode the chain information, a binary feature indicating if two residues associated with an edge are two adjacent (consecutive) residues in the same chain is used. In addition, an edgewise positional encoding (Morehead *et al*. (2021b)) is used for each edge.

#### Node and edge embeddings

After receiving a protein complex graph 𝒢 as input, DProQA applies initial node and edge embedding modules to each node and edge, respectively. We define such embedding modules as *φ*_*h*_ and *φ*_*e*_ respectively, where each *φ* function is represented as a shallow neural network consisting of a linear layer, batch normalization, and LeakyReLU activation function (Xu *et al*. (2015)). Such node and edge embeddings are then fed as an updated input graph to the Gated Graph Transformer.

### 3.2 Gated graph transformer architecture

Unlike other graph neural network (GNN)-based structure scoring methods (Wang *et al*. (2021); Han *et al*. (2021)), which define edges using a fixed distance threshold so that each graph node may have a different number of incoming and outgoing edges, DProQA constructs and operates on k-NN graphs where all nodes are connected to the same number of neighbors. However, in the context of k-NN graphs, each neighbor’s information is, by default, given equal priority during information updates. Here, we may desire to imbue our graph neural network with the ability to automatically set the priority of different nodes and edges during the graph message passing. Consequently, we design GGT, a gated neighborhood-modulating graph transformer inspired by Veličković *et al*. (2017); Dwivedi and Bresson (2020); Morehead *et al*. (2021b) to update the features of the nodes and edges. Formally, to update the network’s node embeddings **h**_*i*_ and edge embeddings **e**_*ij*_, we define a single layer of the GGT as:

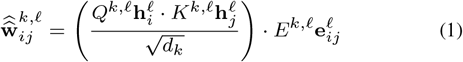

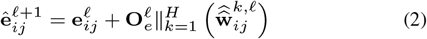

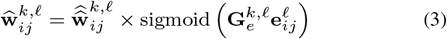

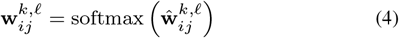

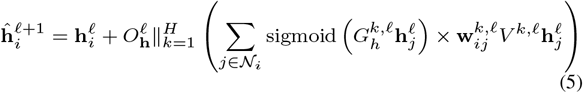

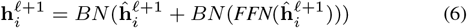

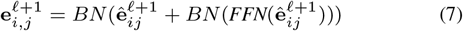

In particular, the GGT adds on top of the standard graph transformer architecture (Dwivedi and Bresson (2020)) two information gates through which the network can modulate node and edge information flow, as shown in Figure 2. Several main operations in Figure 2 are described by the following equations. Equation (1) computes the intermediate attention coefficient 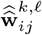 for node pair *i* and *j* in the graph. *Q*^*k,ℓ*^ and *K*^*k,ℓ*^ are learnable parameters, while 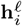 and 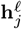 are the node feature vectors at layer *ℓ*. The dot product measures the similarity between the two nodes, and it is normalized by 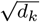, where *d*_*k*_ is the dimension of the attention head. Finally, the result is element-wise multiplied with 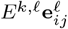, where 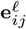 is the edge feature vector and *E*^*k,ℓ*^ is a learnable parameter. Equation (2) updates the intermediate edge feature vector 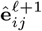 at layer *ℓ* + 1. 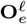 is a learnable parameter, and the concatenation operation 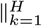 combines the intermediate attention coefficients 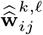 from each attention head *k*. The original edge feature vector 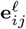 is added to this linear combination via a residual connection. Equation (3) computes the attention coefficients 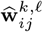 by multiplying the intermediate attention coefficients 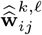 with the edge gate. The edge gate is a sigmoid activation of a linear transformation of the edge features, where 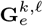 is a learnable parameter. Equation (4) applies the softmax function to normalize the attention coefficients 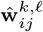, resulting in the final attention weights 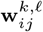. Equation (5) updates the intermediate node feature vector 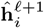 at layer *ℓ* + 1. The concatenation operation 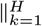 combines the updated node features from each attention head *k*. Specifically, for each attention head *k*, the attention coefficients 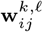 are multiplied by the weight matrix *V* ^*k,ℓ*^ and the feature vector 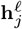 of the neighboring node *j*. These values are then multiplied with a sigmoid output of the gated features of the neighboring node *j*, after which all these values are summed up over the neighbors of node *i*. 𝒩_*i*_ denotes the set of sneighbors of node *i* including itself. The resulting vector is transformed by a learnable matrix 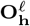. The raw node feature vector 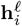 is added to this linear combination via a residual connection. Equation (6) updates the node *i* feature vector at level *ℓ* + **1**, i.e., 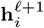. It is obtained by applying batch normalization (BN) to sum of the intermediate node feature vector 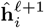 and the output of the feed-forward network (FFN) with BN applied to the 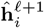. Equation (7) updates 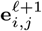 with the same logic as Equation (6). The FFN uses the same structure as described in Dwivedi and Bresson (2020).

**Fig. 2.**
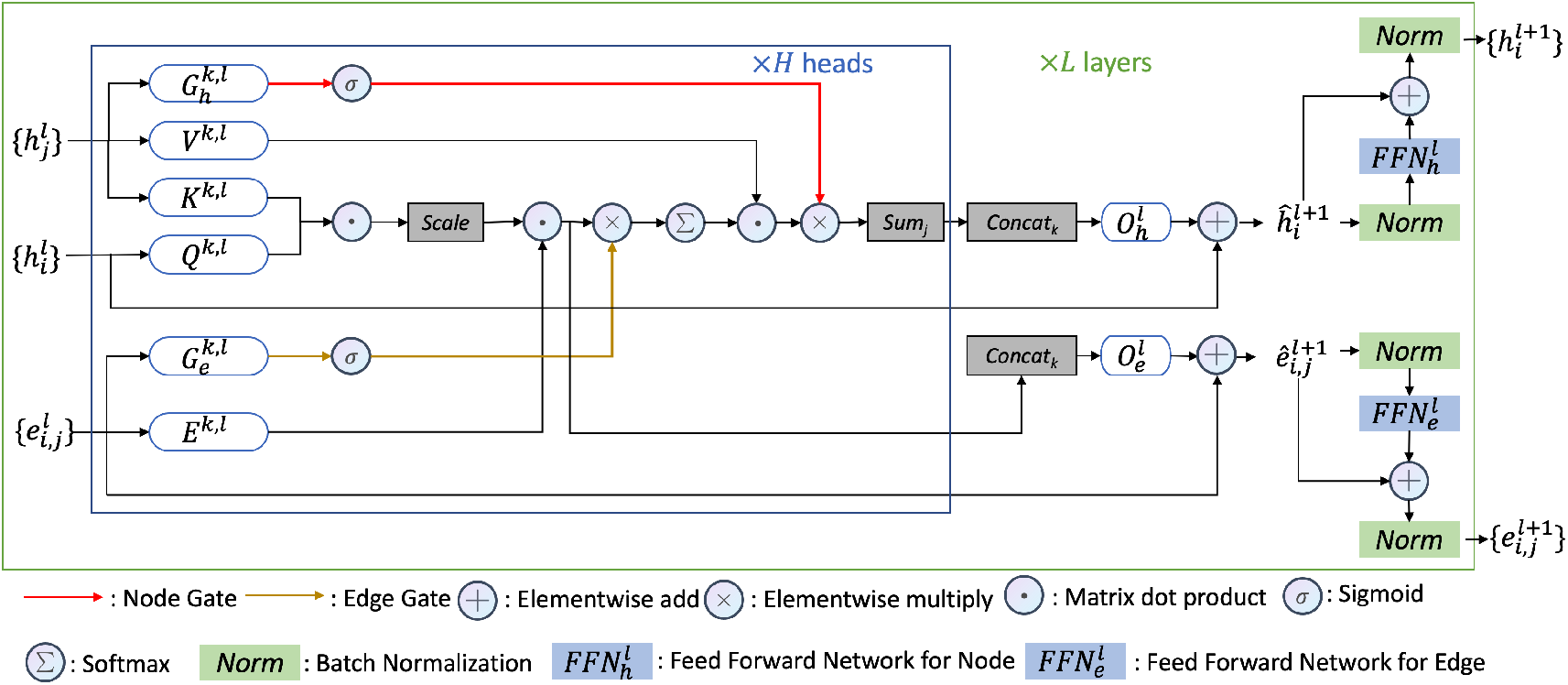
The gated graph transformer (GGT) model architecture for updating the features of nodes of a protein complex graph.

### 3.3 Multi-task graph property prediction

To obtain graph-level predictions for each input protein complex graph, we apply a graph sum-pooling operator on 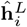 to get the graph embedding **p**. This graph embedding **p** is then fed as input to DProQA’s two read-out modules, where each read-out module consists of a series of linear layers, batch normalization layers, LeakyReLU activation, and dropout layers (Hinton *et al*. (2012)), respectively. The output from the read-out modules is used by a softmax function in the classification output layer to classify the input into the four different quality classes (i.e., Incorrect, Acceptable, Medium, and High). The output from the read-out modules is also used by the regression output layer with a sigmoid activation function to **y** to obtain a single scalar output representing the predicted DockQ score (Basu and Wallner (2016a)) for the protein complex input.

#### Structural quality score prediction loss

To train DProQA’s graph regression head, we used the mean squared error loss 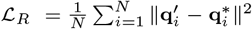. Here, 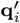 is the model’s predicted DockQ score for example *i*, 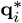 is the ground truth DockQ score for example *i*, and *N* represents the number of examples in a given mini-batch.

#### Structural quality classification loss

To train DProQA’s graph classification head,(we used the) cross-entropy loss 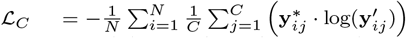. Here, 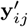 is the predicted probability of the model’s quality belonging to class *j* for example *i*, and 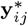 is the ground truth DockQ quality class *j* (e.g., Incorrect) for example *i. N* denotes the number of examples in a given mini-batch and *C* for the number of classes.

#### Overall loss

DProQA’s overall loss is the weighted sum of the two losses above: 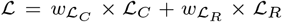. The weights for each constituent loss (e.g.,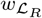) were determined either by performing a grid search or using a lowest-validation-loss criterion for parameter selection. In this project, we set 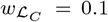 and 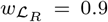. Supplementary Section Implementation and Training Details and Table S1 describe how we implemented, trained, and tuned the DProQA.

### 3.4 Training and test data

#### Multimer-AF2 training dataset

Similar to Morehead *et al*. (2021a), we created a new Multimer-AF2 (MAF2) dataset comprised of multimeric structures predicted by AlphaFold 2 (Jumper *et al*. (2021)) and AlphaFold-Multimer (Evans *et al*. (2021)) structure prediction pipeline on the Summit supercomputer (Gao *et al*. (2021, 2022)). The protein multimer targets for which we predicted structures were obtained from the EVCoupling (Hopf *et al*. (2019)) and DeepHomo (Yan and Huang (2021)) datasets, which consist of both heteromers and homomers. In summary, the MAF2 dataset contains a total of 9, 251 decoys. According to DockQ scores, 20.44% of them are of Incorrect quality, 14.34% of them are of Acceptable quality, 30.00% of them are of Medium quality, and the remaining 35.22% of them are of High quality.

#### Docking Decoy Set for training

The Docking Decoy dataset (Kundrotas *et al*. (2018)), contains 58 protein complex targets. Each target includes approximately 100 incorrect decoys and at least one *near-native* decoy.

The Docking Decoy set and MAF2 set were used together as the training data to train and validate the DProQA, which together include 12, 040 decoys in total. To split the data into training and validation sets, we applied MMseq2 (Mirdita *et al*. (2021)) to cluster all targets’ sequences with 30% sequence identity. Then we selected 70% of the clusters’ decoys as the training set and the rest as the validation set. The combined training set contains 8, 733 decoys, and the validation set contains 3, 407 decoys.

#### Docking Benchmark5.5 AF2 test dataset

The Docking Benchmark 5.5 AF2 (DBM55-AF2) dataset is the first test dataset. We applied AlphaFold-Multimer (Evans *et al*. (2021)) to predict the structures of Docking Benchmark 5.5 targets (Vreven *et al*. (2015)). To avoid the overestimation of the performance of DProQA, we performed 30% sequence identity filtering w.r.t the training and validation data to remove similar targets with >30% sequence identity. Overall, this test dataset contains a total of 15 protein targets with 449 total decoy models, 50.78% of these decoys are of Incorrect quality, 16.70% of them are of Acceptable quality, 30.73% of them are of Medium quality, and the remaining 1.78% of them are of High quality.

More details about MAF2 and DBM55-AF2 generation and how we conducted sequence filtering and selected targets as the blind test set can be found in Supplementary Section Addition Dataset Information.

#### CASP15 EMA experiment

We blindly tested DProQA (Group name: MULTICOM_egnn, ID: 120) in 2022 CASP15 EMA category. DProQA was evaluated on 36 protein complex targets whose exprimental structures were available for us. Each target contains around 350 models from different CASP15 protein quaternary structure predictors.

#### Training labels

DProQA performs two learning tasks simultaneously. In the regression task, DProQA treats true DockQ scores (Basu and Wallner (2016a)) of decoys as its labels. As introduced earlier, DockQ scores are continuous values in the range of [0, 1]. A higher DockQ score indicates a higher-quality structure. In the classification task, DProQA predicts the probabilities that the structure of an input protein complex falls into the Incorrect, Acceptable, Medium, or High-quality category. Labeling a decoy into such quality categories was made according to its true DockQ score.

The true DockQ scores of the models in MAF2 and DBM55-AF2 were calculated by using the DockQ tool (Basu and Wallner (2016a)) to compare them with their corresponding true structures. The Docking Decoy Set provides interface root mean squared deviations (iRMSDs), ligand RMSDs (LRMSs), and fractions of native contacts (*f*_*nat*_) for each decoy. We directly used Equation 1 and 2 in Basu and Wallner (2016a) to convert these scores to DockQ scores. The DockQ scores were then converted into four discrete categories: Incorrect, Acceptable, Medium, or High quality according to Basu and Wallner (2016a).

### 3.5 Evaluation setting

#### Baseline methods

We compared DProQA with three typical methods: ZRANK2, GOAP and GNN_DOVE. ZRANK2 is a method using a linear weight scoring function for evaluating protein complex structures. GOAP score is composed of all-atoms level distance-dependent and orientation-dependent potentials. GNN DOVE is an atom-level graph attention-based method for protein complex structure evaluation. It extracts the interface areas of a protein complex structure to build its input graph.

#### DProQA variants

Besides the standard DProQA model, we also report results on the DBM55-AF2 dataset for a selection of DProQA variants curated in this study. The DProQA variants includes DProQA_GT which employs the original Graph Transformer architecture Dwivedi and Bresson (2020); DProQA_GTE which employs the GGT with only its edge gate enabled; and DProQA_GTN which employs the GGT with only its node gate enabled.

#### Evaluation metrics

We evaluated the methods using two main metrics. The first metric measures how many qualified decoys are found within a model’s predicted Top-*N* structure ranking for a target. Within this framework, a method’s overall hit (success) rate is defined as the number of protein complex targets for which it ranks at least one acceptable, medium or higher-quality decoy within its Top-*N* ranked decoys, which is a metric used by the Critical Assessment of Protein-Protein Interaction (CAPRI) (Lensink and Wodak (2014)). In this work, we report the methods’ Top-10 hit rates. A hit rate is represented by three numbers separated by the character /. These three numbers, in order, represent how many decoys with Acceptable or higher-quality, Medium or higher-quality, and High quality are among the Top-N ranked decoys. The second metric measures the ranking loss for each method. Here, the per-target ranking loss is defined as the difference between the DockQ score of a target’s best decoy and the DockQ score of the top decoy selected by the ranking method. As such, a lower ranking loss indicates a better ranking ability.

## 4 Results

### 4.1 Performance on the DBM55-AF2 dataset

Table 2 presents the ranking loss for all methods on the DBM55-AF2 dataset. DProQA achieves the best ranking loss of 0.049 which is 86.56% lower than ZRANK2’s ranking loss 0.372, 60.16% lower than GOAP’s ranking loss of 0.123 and 84.19% lower than GNN_DOVE’s ranking loss of 0.31. Furthermore, for 4 targets, DProQA correctly selects the Top-1 model and achieves 0 ranking loss. Additionally, DProQA and GOAP achieve the lowest loss on 5 targets, while ZRANK2 gets the lowest loss on 2 targets and GNN_DOVE on 2 targets. Notably, DProQA_GT, DProQA_GTE, and DProQA_GTN’s losses are also lower than the three baseline methods, but they are higher than that of DProQA.

**Table 2.** The DockQ score ranking loss on the DBM55-AF2 dataset. DProQA denotes the final Gated Graph Transformer, DProQA_GT denotes the original Graph Transformer architecture, DProQA_GTE denotes the GGT with only its edge gate enabled, and DProQA_GTN denotes the GGT with only its node gate enabled. The final row reports the mean and standard deviation (Std) of the ranking loss of the different methods.

| Target | DProQA | DProQA_GT | DProQA_GTE | DProQA_GTN | ZRANK2 | GOAP | GNN_DOVE |
| --- | --- | --- | --- | --- | --- | --- | --- |
| 6ALO | 0.0 | 0.156 | 0.156 | 0.0 | 0.345 | 0.331 | 0.382 |
| 3SE8 | 0.079 | 0.041 | 0.041 | 0.079 | 0.735 | 0.0 | 0.408 |
| 5GRJ | 0.024 | 0.012 | 0.095 | 0.012 | 0.774 | 0.23 | 0.595 |
| 6A77 | 0.037 | 0.062 | 0.0 | 0.037 | 0.583 | 0.59 | 0.589 |
| 4M5Z | 0.015 | 0.026 | 0.026 | 0.015 | 0.221 | 0.133 | 0.269 |
| 4ETQ | 0.0 | 0.76 | 0.0 | 0.748 | 0.759 | 0.0 | 0.748 |
| 5CBA | 0.052 | 0.038 | 0.052 | 0.058 | 0.047 | 0.007 | 0.047 |
| 5WK3 | 0.114 | 0.114 | 0.114 | 0.186 | 0.0 | 0.109 | 0.109 |
| 5Y9J | 0.0 | 0.0 | 0.0 | 0.0 | 0.202 | 0.0 | 0.423 |
| 6BOS | 0.081 | 0.081 | 0.0 | 0.0 | 0.087 | 0.09 | 0.053 |
| 5HGG | 0.051 | 0.051 | 0.121 | 0.051 | 0.051 | 0.051 | 0.047 |
| 6A0Z | 0.207 | 0.207 | 0.207 | 0.207 | 0.218 | 0.214 | 0.206 |
| 3U7Y | 0.0 | 0.021 | 0.0 | 0.0 | 0.772 | 0.0 | 0.021 |
| 3WD5 | 0.011 | 0.011 | 0.011 | 0.0 | 0.704 | 0.011 | 0.666 |
| 5KOV | 0.065 | 0.08 | 0.085 | 0.087 | 0.008 | 0.078 | 0.083 |
| Mean ± Std | **0.049 ± 0.054** | 0.111 ± 0.182 | 0.061 ± 0.064 | 0.099 ± 0.185 | 0.372 ± 0.3 | 0.123 ± 0.158 | 0.310 ± 0.245 |

Table 3 summarizes all the methods’ hit rates on the DBM55-AF2 dataset, which contains 15 targets. Notably, DProQA excels in achieving the highest hit rate for ranking medium-quality decoys of all the 10 targets that have at least one medium- or high-quality decoy. In terms of selecting high-quality decoys, DProQA, ZRANK2, and GOAP effectively identify high-quality decoys for all the 3 targets that have high-quality decoys. However, it is ranked behind ZRANK2 and GOAP in selecting acceptable-quality decoys, i.e., it is able to rank at least one acceptable quality model in the top 10 for 12 out of 15 targets that have at least one acceptable decoy, one fewer than ZRANK2 and GOAP.

**Table 3.** Per-target and overall hit rates on the DBM55-AF2 dataset. The last column represents each target’s best-possible Top-10 result, which is an upper limit of the hit rates. The ‘a/b/c’ values for each target represent the number of top-10 ranked decoys that have the Acceptable or higher-quality, Medium or higher-quality, and High-quality, respectively. The ‘a/b/c’ values in the Summary row reports the number of targets for which each method successfully ranks at least one decoy of the Acceptable or higher-quality, Medium or higher-quality, and High-quality within top 10, respectively

| Target | DProQA | DProQA_GT | DProQA_GTE | DProQA_GTN | ZRANK2 | GOAP | GNN_DOVE | BEST |
| --- | --- | --- | --- | --- | --- | --- | --- | --- |
| 6ALO | 9/2/0 | 10/0/0 | 10/0/0 | 10/2/0 | 9/0/0 | 9/0/0 | 6/0/0 | 10/2/0 |
| 3SE8 | 8/8/0 | 9/9/0 | 8/8/0 | 8/8/0 | 2/2/0 | 8/8/0 | 4/3/0 | 10/10/0 |
| 5GRJ | 10/10/0 | 9/9/0 | 10/10/0 | 9/9/0 | 5/4/0 | 10/9/0 | 6/6/0 | 10/10/0 |
| 6A77 | 7/7/0 | 7/7/0 | 8/8/0 | 8/8/0 | 4/4/0 | 3/3/0 | 3/3/0 | 8/8/0 |
| 4M5Z | 10/10/1 | 10/10/0 | 10/10/0 | 10/10/0 | 10/10/1 | 10/10/1 | 10/10/0 | 10/10/1 |
| 4ETQ | 1/1/0 | 1/1/0 | 1/1/0 | 1/1/0 | 1/1/0 | 1/1/0 | 0/0/0 | 1/1/0 |
| 5CBA | 10/10/1 | 10/10/0 | 10/10/0 | 10/10/1 | 10/10/4 | 10/10/2 | 10/10/3 | 10/10/6 |
| 5WK3 | 0/0/0 | 0/0/0 | 0/0/0 | 0/0/0 | 3/0/0 | 0/0/0 | 0/0/0 | 3/0/0 |
| 5Y9J | 4/0/0 | 6/0/0 | 5/0/0 | 4/0/0 | 5/0/0 | 5/0/0 | 2/0/0 | 8/0/0 |
| 6BOS | 10/10/0 | 10/10/0 | 10/10/0 | 10/10/0 | 10/10/0 | 10/10/0 | 10/10/0 | 10/10/0 |
| 5HGG | 8/0/0 | 8/0/0 | 8/0/0 | 8/0/0 | 10/0/0 | 10/0/0 | 10/0/0 | 10/0/0 |
| 6A0Z | 0/0/0 | 0/0/0 | 0/0/0 | 0/0/0 | 0/0/0 | 0/0/0 | 1/0/0 | 3/0/0 |
| 3U7Y | 2/2/1 | 2/2/1 | 2/2/1 | 2/1/0 | 1/1/1 | 2/2/1 | 2/2/1 | 2/2/1 |
| 3WD5 | 10/8/0 | 9/8/0 | 9/8/0 | 9/8/0 | 6/4/0 | 10/8/0 | 8/6/0 | 10/10/0 |
| 5KOV | 0/0/0 | 0/0/0 | 0/0/0 | 0/0/0 | 0/0/0 | 1/0/0 | 0/0/0 | 2/0/0 |
| Summary | **12/10/3** | 12/9/1 | 12/9/1 | 12/10/1 | **13/9/3** | **13/9/3** | 12/8/2 | 15/10/3 |

### 4.2 Impact of node and edge gates

The results in Tables 2 and 3 show that using both edge and node gates (DProQA) performs better than using only the edge gates (DProQA_GTE) or the node gates (DProQA_GTN), whose accuracy is better than or equal to not using any gate (DProQA_GT). Therefore, this ablation study specifically demonstrates that the edge and node gates are useful for predicting the quality of protein complex structures.

### 4.3 Performance in 2022 CASP15 EMA experiment

DProQA (team: MULTICOM_egnn) participated in the Estimation Model Accuracy(EMA) category of the CASP15 running from May to August 2022. We collected all EMA prediction results from CASP15 website CASP15 (2022) and used US-align Zhang *et al*. (2022) to calculate the TM-score Zhang and Skolnick (2004) ranking loss for all 36 targets whose true structures were available for us to perform evaluation. Figure 3 reports all CASP15 single-model EMA methods’ average TM-score ranking loss, where DProQA ranked **3rd**. DProQA achieved a 0.200 ranking loss. All single-model methods’ average ranking loss is 0.307. The result of TM-score ranking loss for all the CASP15 multi-model and single-model EMA methods can be found in the supplementary Figure S3. DProQA performed even better than 3 out of 9 multi-model EMA methods.

**Fig. 3.**
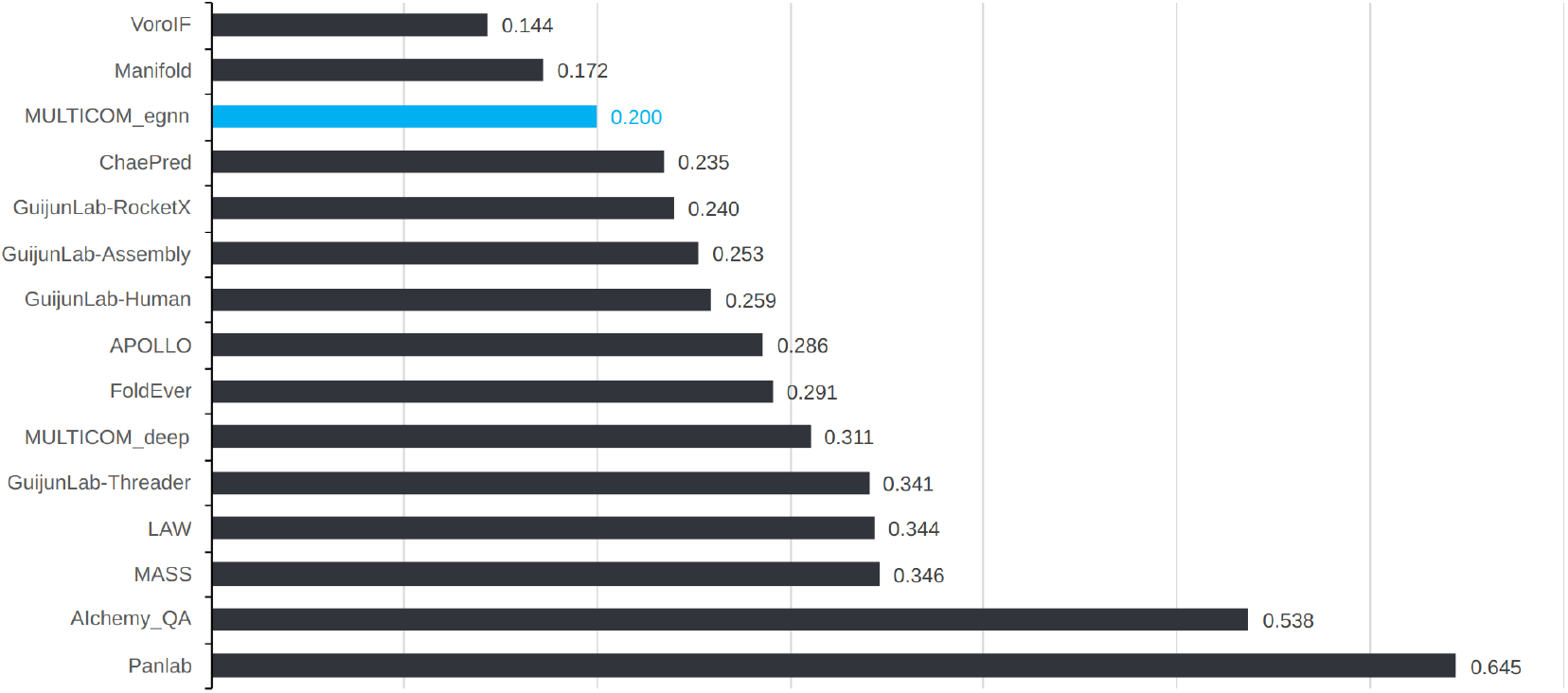
The average TM-score ranking loss for all single-model methods. MULTICOM_egnn ranked 3rd among all single-model methods

Figure 4 illustrates the distribution of the MULTICOM_egnn’s ranking loss on the 36 CASP15 EMA targets. The vertical dashed black line is the mark for the mean value. 23 out of 36 data points are located on the left side of the black line. This loss distribution is right-skewed, where the skewness value is 0.751.

**Fig. 4.**
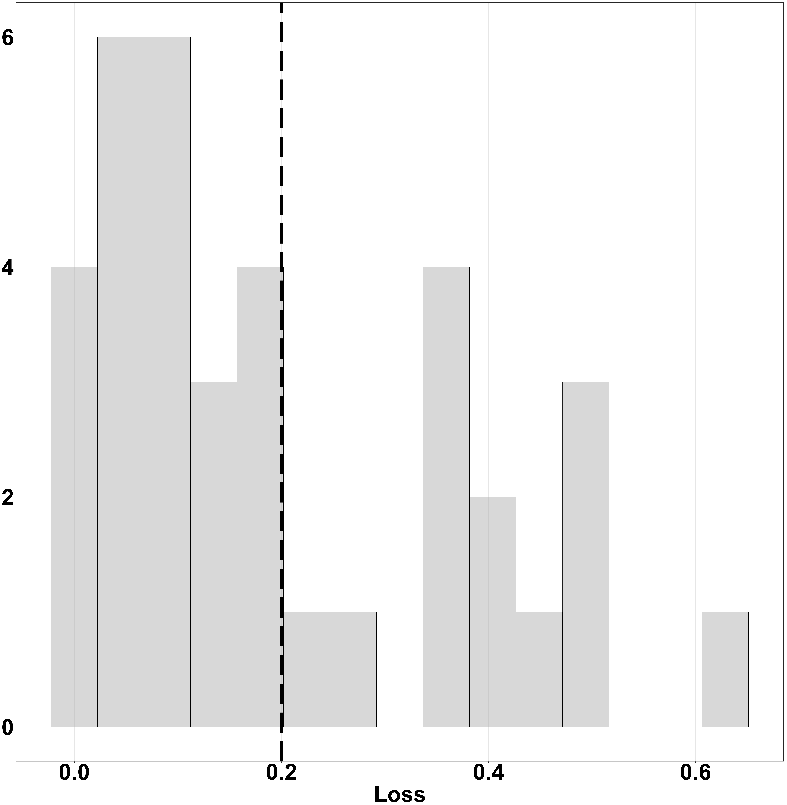
MULTICOM_egnn’s loss histogram for CASP15 targets. The black dashed vertical line represents the position of the mean value.

Figure 5 shows that MULTICOM_egnn successfully selected a high-quality model with a very low ranking loss (i.e., 0.0014) for target H1111 (PDB code: 7QIJ) which is a Hetero 27-mer with a sequence length of 8460. Figure 5(a) is the histogram of all server methods’ model TM-scores for target H1111. Most models’ TM-scores are low, yet still some are high-quality prediction models. In Figure 5(b), from left to right, the three protein complex structures shown are the corresponding native structure, the true TOP-1 model, and the MULTICOM_egnn top selected model, respectively. The top model selected by MULTICOM_egnn has a high TM-score of 0.9816. The ranking loss of MULTICOM_egnn for this target is the lowest among all the CASP15 EMA methods.

**Fig. 5.**
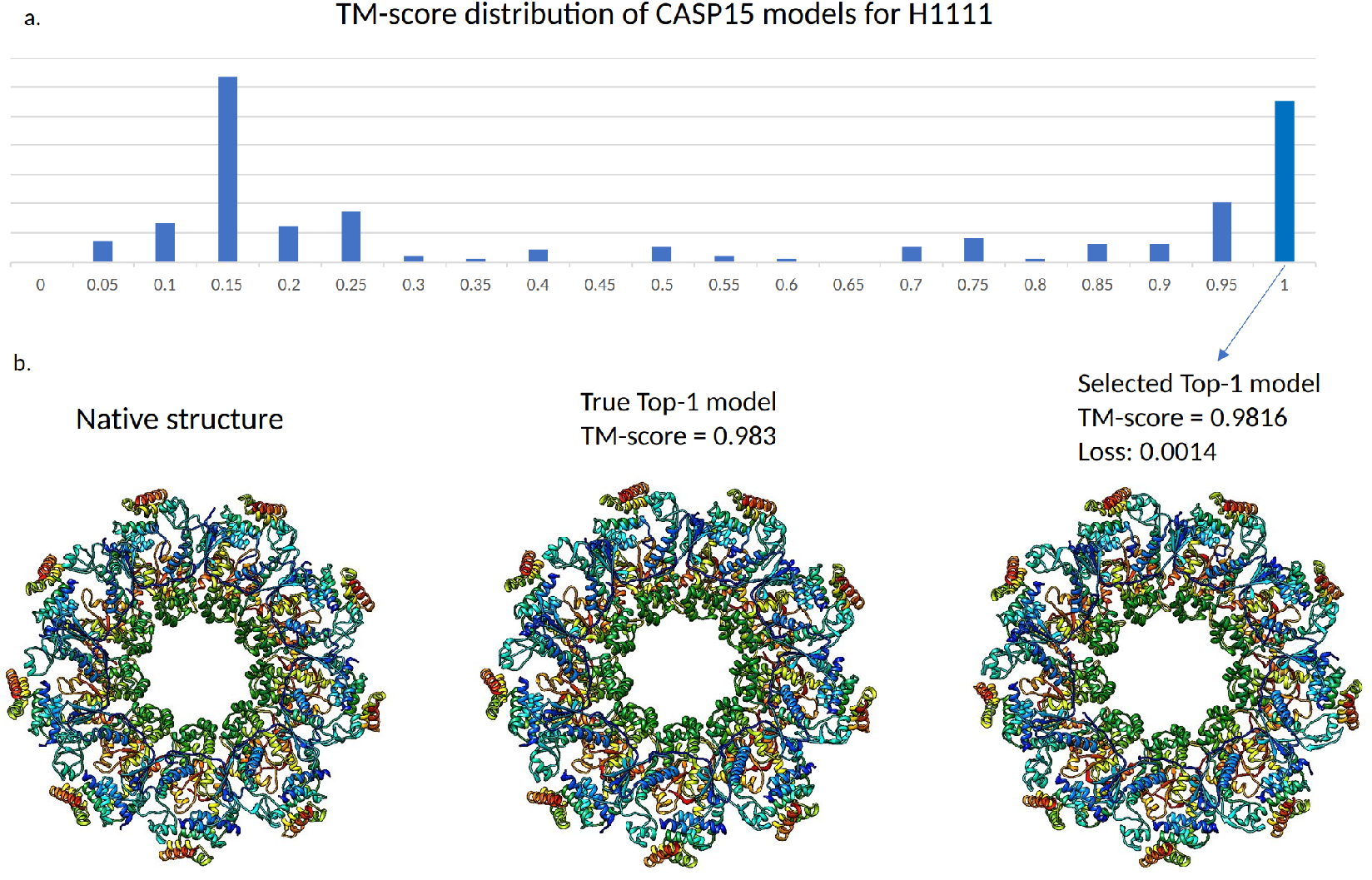
Target H1111. (A) TM-score distribution of CASP15 models of H1111. (B). From left to right, the three protein complex structures shown are the corresponding native structure, the true TOP-1 model, and the MULTICOM_egnn top selected model, respectively. Here, MULTICOM_egnn achieved a 0.0014 TM-score ranking loss.

## 5 Discussion

Compared to the general high accuracy of predicted tertiary structures (Jumper *et al*. (2021)), the average accuracy of quaternary structures predicted for protein complexes (multimers) is still relatively low (Bryant *et al*. (2022)), making the selection of good models from a pool of decoys harder. We observe this not only in some popular complex datasets Lensink and Wodak (2014); Kundrotas *et al*. (2018), but also in our newly-built DBM55-AF2 sets. For instance, some targets like 6A0Z and 3U7Y in the DBM55-AF2 set only have a few decoys with acceptable or higher quality, while the rest of decoys have very low quality. If no model of acceptable or higher quality is ranked at the top, the ranking loss will be very high (see some examples in Table 2). Therefore, a significant challenge in ranking the models of protein multimers is to identify a few good models in a large pool of mostly bad models.

A more common way to evaluate the ranking ability of the quality assessment (QA) methods for protein complexes is the hit rate, a standard method used by CAPRI. However, a hit rate only measures the number of qualified decoys in the TOP N ranked models, without measuring the difference between the best possible model and the top-ranked model. Therefore, in this work, we also apply the loss metric widely used in evaluating the quality assessment methods for protein tertiary structures to the quality assessment for protein quaternary structures.

Considering these two metrics together helps us evaluate a protein multimeric QA method’s ranking ability more effectively. For example, ZRANK2 which has slightly better hit-rate performance than DProQA on the DBM55-AF2 set, while its loss is much higher than DProQA’s loss. Overall, DProQA demonstrates consistently good performance on our internal benchmark as well as the most rigorous blind CASP15 benchmark.

On the DMB55-AF2 test dataset, DProQ’s average running time of assessing the quality of the models of each target is about 12 seconds, which is much faster than GOAP and GNN_DOVE but slower than ZRANK2-an energy-based method (see supplementary Table S4 for the detailed execution time of the four EMA methods).

It should be emphasized that AlphaFold-multimer has the capability to assess the quality of a structural model by utilizing its own ipTM score. Nevertheless, the ipTM score is heavily influenced by the evolutionary information (e.g., multiple sequence alignments) and templates. In contrast, DProQA’s prediction is solely based on a single 3D model and therefore provides a fast and complementary estimation of the quality of a structural model.

## 6 Conclusion

In this work, we present DProQA - a gated graph transformer for protein complex structure assessment. Our rigorous experiments and CASP15 results demonstrate that DProQA performs relatively well in ranking decoy models of protein complexes and the gated message passing in the transformer is useful for improving its performance. Both the tool and the new datasets consisting of multimer models predicted by AlphaFold2 and AlphaFold-Multimer are made publicly available for the community to further advance the field.

