## Supplementary for "A Gated Graph Transformer for Protein Complex Structure Quality Assessment and its Performance in CASP15"

### Implementation and Training Details

DProQA was implemented on top of PyTorch [1], PyTorch Lightning [3], and the Deep Graph Library [4]. We trained the deep learning models by using the ADAMW optimizer [5] and performed early stopping with a patience of 15 epochs. Also, we applied Stochastic Weight Averaging (SWA) [2] to smooth the training loss. We choose the best model based on the validation set's performance. All parameters' settings are described in Table S1. The training config file is: https://github.com/jianlincheng/DProQA/blob/main/configs/Lab_saw_mulitase_gate_af2_decoy_knn10_seed2222.json

DProQA was trained and tested on the platform with following software and hardware:

System: Ubuntu 22.04 (LTS)
CPU: Intel(R) Core (TM) i9-9900K CPU @ 3.60GHz.
GPU: GeForce RTX 2080 SUPER (2X).
RAM: 64 GB.

**Table S1.** Hyperparameter values searched to obtain the good performance on the validation data. The final parameter values for the standard DProQA model are denoted in **bold**.

| Hyperparameter | Search Space |
|---|---|
| Weight of $L_C$ ($w_{LC}$) | 0.1 |
| Weight of $L_R$ ($w_{LR}$) | **0.9**, 0.8, 0.7, 0.6, 0.5 |
| Number of GGT Layers | 1, **2**, 3, 4, 6 |
| GGT Dropout Rate | 0.1, 0.2, 0.3, **0.4**, 0.5 |
| Read-Out Module Dropout Rate | 0.1, 0.2, 0.3, 0.4, **0.5** |
| Number of Attention Heads | 8 |
| Hidden Dimension | 32, **64**, 128 |
| Non-Linearities | LeakyReLU |
| Learning Rate | 0.001, **0.005**, 0.01, 0.05, 0.1 |
| Weight Decay Rate | 0.001, **0.002**, 0.01, 0.02 |

| | |
|---|---|
| Normalization | LayerNorm, **BatchNorm** |
| Graph Pooling Operator | Mean, Max, **Sum** |

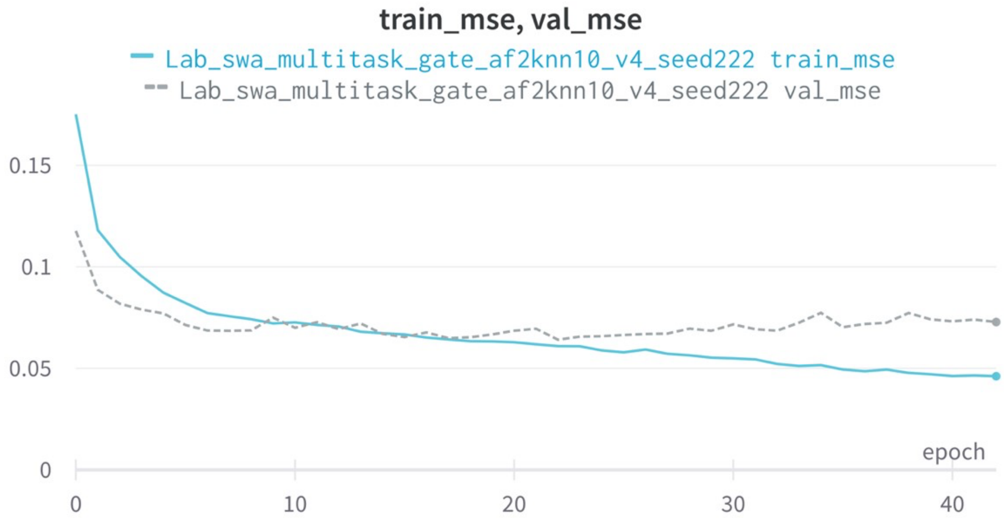


Figure S1: Mean squared error (MSE) loss curve on the training and validation dataset. The solid blue line is training loss, and the dashed grey line is validation loss.

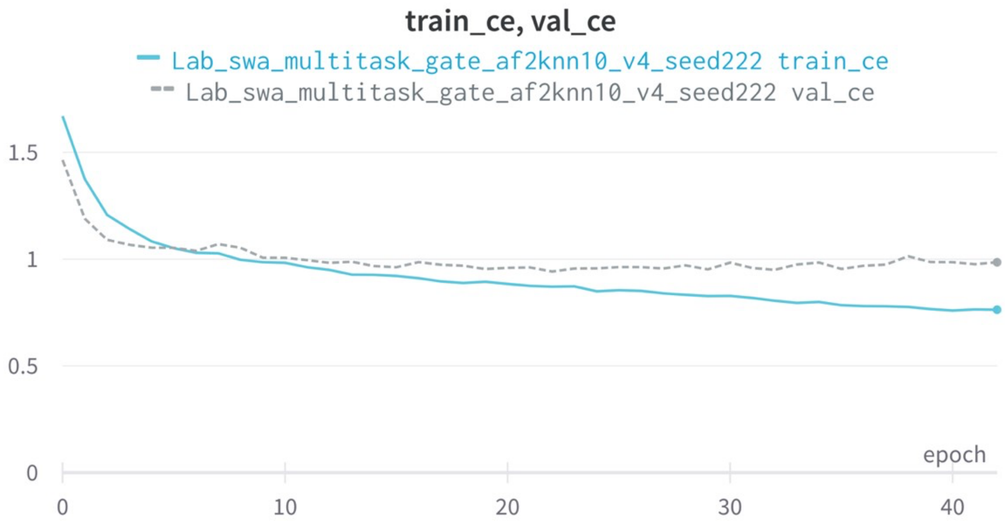


Figure S2: Cross entropy loss curve on the training and validation dataset. The solid blue line is the training loss, and the dashed grey line is the validation loss.

### Additional Data Information

**Multimer-AF2 dataset:** For the structures we predicted using AlphaFold2 [6], before structure prediction, we inserted 20-glycine residues in between chains in the multimeric input sequences to facilitate multimeric structure generation with AlphaFold2. For structures predicted with AlphaFold-Multimer, we directly input the original multimeric sequences into the model's prediction pipeline. Only the top 1 model ranked by AlphaFold2 [6] or AlphaFold-Multimer [7] was selected to be included into the training dataset. Table S2 is the statistics of the MAF2 dataset.

**Table S2.** Summary of the MAF2 dataset.

| | Incorrect | Acceptable | Medium | High | Mean DockQ | Median DockQ |
|---|---|---|---|---|---|---|
| AlphaFold-Multimer | 590 | 371 | 1237 | 1878 | 0.66 | 0.78 |
| AlphaFold 2 | 1301 | 956 | 301 | 1380 | 0.52 | 0.67 |
| Total | 1891 | 1327 | 2775 | 3258 | 0.59 | 0.74 |

**Docking Benchmark 5.5 AF2 test dataset:** We used AlphaFold-Multimer [7] to predict the structures of Docking Benchmark 5.5 targets [8]. Specifically, this test dataset consists of five randomly sampled targets from each of the Benchmark 5.5 dataset's three difficulty categories (i.e., Rigid-Body, Medium, Difficult). After obtaining the targets for this test dataset, we calculated the DockQ score [9] for each decoy model for the targets and then filtered out the targets that do not contain any decoy of Acceptable or higher quality. Moreover, we performed 30% sequence by MMseq2 [10] identity filtering w.r.t the training and validation data to remove similar targets. After the filtering, this test dataset contains 15 protein targets with 449 decoy models associated with them. Table S3 shows the quality of the decoys for each target in the test dataset.

**Table S3.** Summary of the quality of the Docking Benchmark 5.5 AF2 dataset.

| Target | Incorrect | Acceptable | Medium | High | Mean DockQ | Median DockQ |
|---|---|---|---|---|---|---|
| 6AL0 | 4 | 24 | 2 | 0 | 0.28 | 0.26 |
| 3SE8 | 15 | 3 | 12 | 0 | 0.33 | 0.19 |
| 5GRJ | 8 | 1 | 21 | 0 | 0.52 | 0.68 |
| 6A77 | 22 | 0 | 8 | 0 | 0.23 | 0.08 |
| 4M5Z | 0 | 0 | 29 | 1 | 0.61 | 0.59 |
| 4ETQ | 29 | 0 | 1 | 0 | 0.08 | 0.04 |
| 5CBA | 0 | 0 | 24 | 6 | 0.78 | 0.78 |
| 5WK3 | 27 | 3 | 0 | 0 | 0.17 | 0.18 |
| 5Y9J | 22 | 8 | 0 | 0 | 0.16 | 0.12 |
| 6B0S | 0 | 0 | 30 | 0 | 0.57 | 0.57 |
| 5HGG | 2 | 27 | 0 | 0 | 0.29 | 0.29 |
| 6A0Z | 27 | 3 | 0 | 0 | 0.1 | 0.08 |
| 3U7Y | 28 | 0 | I | 1 | 0.1 | 0.04 |
| 3WD5 | 16 | 4 | 10 | 0 | 0.29 | 0.18 |

| | | | | | | |
|---|---|---|---|---|---|---|
| 5KOV | 28 | 2 | 0 | 0 | 0.19 | 0.19 |
| Total | 228 | 75 | 138 | 8 | 0.32 | 0.19 |

### Additional results on DMB55-AF2 test dataset

**Table S4** reports the running time of DProQA, GOAP, ZRANK2, and GNN_DOVE on the DBM55-AF2 test set. ZRANK2 is a physical energy score-based method with an average running time of 0.748s, which is faster than the other three methods. DProQA is the second-fastest method, with an average time of 11.897s, which is two times faster than the statistical-based method GOAP and almost seven times faster than the deep learning-based method GNN-DOVE. DProQA and GNN_DOVE are GPU-based methods. One GeForce RTX 2080 SUPER was used by them for inference if possible. As GNN_DOVE requires large memory for some large targets, a more advanced GPU – V100 with 32GB was used by GNN_DOVE for the long sequence targets.

**Table S1**. The execution time of the four methods (seconds) on DMB55-AF2 teste dataset. The last row is average running time on the 15 targets. *: the V100 (32GB) was used by GNN_DOVE to estimate model accuracy for the target because the GeForce RTX 2080 SUPER did not have sufficient memory.

| Target | DProQA | GOAP | ZRANK2 | GNN_DOVE |
|---|---|---|---|---|
| 6AL0 | 11.841 | 25.001 | 0.849 | 135.32 |
| 3SE8 | 13.761 | 34.787 | 0.849 | 270.925* |
| 5GRJ | 9.943 | 19.317 | 0.276 | 64.858 |
| 6A77 | 10.755 | 21.988 | 0.886 | 104.534 |
| 4M5Z | 11.933 | 27.759 | 0.87 | 146.679 |
| 4ETQ | 12.175 | 29.795 | 0.862 | 227.961* |
| 5CBA | 8.949 | 14.56 | 0.308 | 39.362 |
| 5WK3 | 10.466 | 1.235 | 0.889 | 100.112 |
| 5Y9J | 15.044 | 42.843 | 0.821 | 299.672* |
| 6B0S | 10.485 | 21.203 | 0.833 | 95.572 |
| 5HGG | 9.23 | 18.004 | 0.626 | 50.429 |
| 6A0Z | 12.737 | 31.106 | 0.891 | 170.141 |
| 3U7Y | 13.644 | 34.511 | 0.942 | 204.413* |
| 3WD5 | 15.251 | 42.731 | 0.844 | 228.51* |
| 5KOV | 12.241 | 32.797 | 0.317 | 104.507 |
| Mean | 11.897 | 26.509 | 0.748 | 149.533 |

### Additional results on the CASP15 dataset

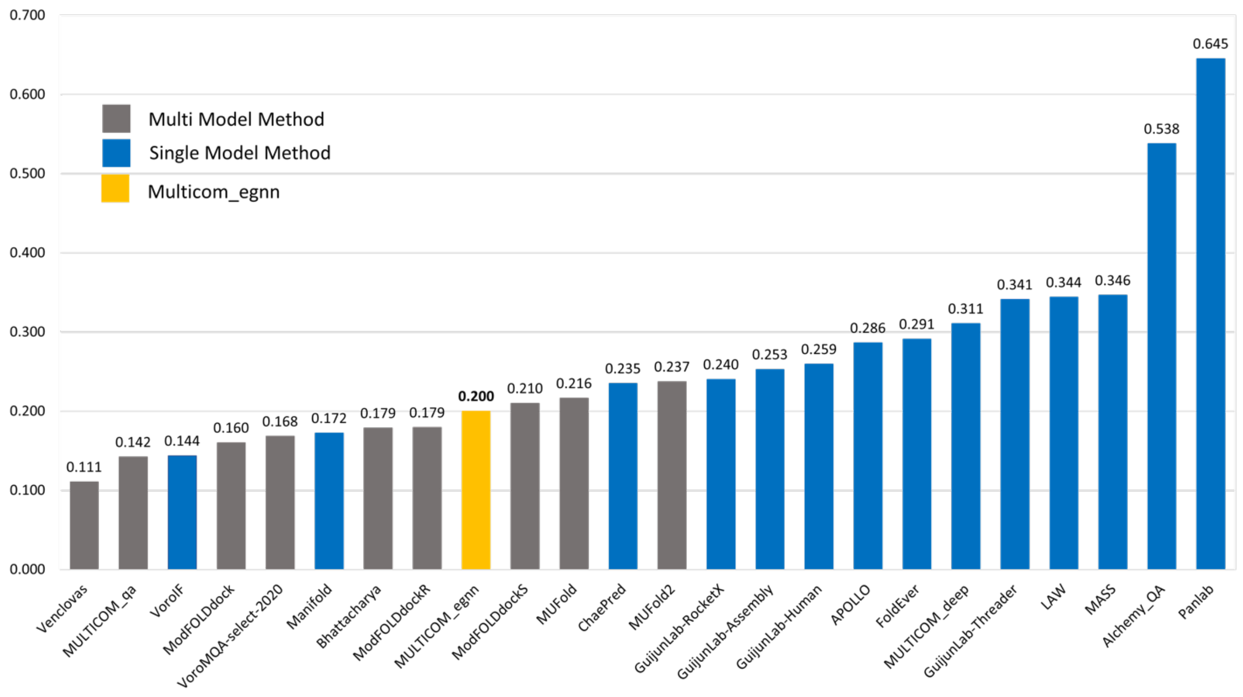


**Figure S3**. The TM-score ranking loss of all CASP15 multi-model and single-model EMA methods on the CASP15 dataset. Grey bars represent multi-model methods and blue bars represent single-model Methods. Yellow bar is Multicom_egnn (i.e., DProQA). DProQA ranks No. 3 among the single-model methods and No. 9 among all the EMA methods. DProQA performs better than three multi-model methods.
